# Disulfiram Exerts anti-pulmonary Fibrosis Effect by Activating PGE2 Synthesis

**DOI:** 10.1101/2023.04.16.537099

**Authors:** Xiaolin Pei, Fangxu Zheng, Yin Li, Zhoujun Lin, Yupeng Zhang, Xiao Han, Ya Feng, Fei Li, Juan Yang, Tianjiao Li, Zhenhuan Tian, Ke Cao, Dunqiang Ren, Chenggang Li

## Abstract

Idiopathic pulmonary fibrosis (IPF) is marked with the replacement of normal alveolar tissue by thicker and harder fibrous material, damaged exchange ability. Currently, nintedanib and pirfenidone, are the only FDA-approved drugs with limited efficacy for IPF, which indicated an urgent need to explore new therapies. Disulfiram (DSF), an acetaldehyde dehydrogenase inhibitor, used as anti-alcohol treatment. Despite reported with anti-hepatic fibrosis effect of DSF, the underlying mechanism remains unclear. In our study, DSF exhibited regulative impact on abnormal proliferation, EMT and ECM production in cell models of IPF including primary DHLF-IPF cells and TGF-β1-stimulated A549 cells. The absence of COX-2 was restored by DSF treatment, together with elevated prostaglandin biosynthesis both in vitro and in vivo models of IPF. Furthermore, the anti-fibrotic effect of DSF was impeded with COX-2 knockdown or pharmacological inhibition in TGF-β1-stimulated A549 cells, however, exogenous PGE2 reclaimed with anti-EMT function. In established animal model of IPF, DSF ameliorated declined lung function and histopathological changes, and restrained the lung hydroxyproline content. Together, these findings suggest that the anti-fibrotic effect of DSF was achieved through re-activation of COX-2 mediated PGE2 biosynthesis. The above results suggest that DSF can be applied therapeutically in fibrotic conditions.

## 1. Introduction

Idiopathic pulmonary fibrosis (IPF) is a chronic, progressive fibrosis interstitial pneumonia, characterized by the excessive accumulation of extracellular matrix and fibrotic tissue in the lungs [1]. The median survival time is about 2-3 years after diagonosis [2]. Clinically, Nintedanib and pirfenidone, two anti-fibrosis agents approved by U.S food and drug administration (FDA), can slow the decline rate of lung function in patients with IPF, but there are certain some side effects and poor prognosis [3]. Though the pathogenesis is not well illustrated, Epithelial-mesenchymal transition (EMT), and ECM deposition are considered major changes in IPF [4]. The morphology and structure of alveolar epithelial cells changed over the process of self-repair after injury associated with EMT and ECM deposition [5]. A number of essential cytokines contribute to EMT in alveolar epithelial cells, and transforming growth factor (TGF-β1) was identified as the key elements for fibrosis [6]. In vitro study, TGF-β1 induces morphological change, extracellular matrix deposition, tight junctions destroys between cells, and gain-of-function with migrate ability in culturing epithelial cells [7].

IPF patients are characterized with down regulated COX-2 expression and its main metabolite prostaglandin E2 (PGE2), which is the terminal product of COX-2 regulation in arachidonic acid metabolic pathway [8]. PGE2 was regarded as an anti-fibrosis gene and showed the contribution on activation of lung fibroblasts and excessive deposition of collagen in TGF-β1-induced COX-2 depression [9]. The differentiation of fibroblasts into myofibroblasts is the fundamental mechanism of the occurrence and development of IPF. The level of PGE2 up-regulation was capable to reverse differentiation phenotype by inhibiting α-SMA and collagen deposition [10]. In addition, PGE2 inhibits the EMT progression by binding to and activating prostaglandin receptors, indicating that the COX-2/PGE2/EPs axis plays a major role in inhibiting EMT [11]. Disulfiram, FDA-approved drug for several decades, is a safe, well-tolerated, inexpensive agent which was supported in alcohol dependence, and it demonstrated the effects of anti-cancer [12], antiviral [13], as well as metabolic dysfunction improvement [14]. DSF down-regulates the level of aldehyde dehydrogenase family 1 (ALDH1) in fibroblasts, thereby preventing mucosal fibrosis in human and mouse eye scar formation [15]. What’s more, DSF prevents renal fibrosis [16] and liver fibrosis [17] via an oxidative mechanism. It’s reported that DSF inhibits EMT to reduce cell metastasis. DSF suppressed the morphological change, EMT-markers expression, cell migration and invasion in TGF-β1-induced EMT of oral squamous cell carcinoma (OSCC) cells [18]. DSF further existed excellent anti-tumor activity after complexing with copper ion, which can dramatically inhibit the EMT, migration and metastasis of breast cancer cells stimulated with TGF-β1 [19]. However, there was no research on DSF treating IPF via regulating COX-2/PGE2 signal axis.

The aim of our study was to ascertain the effect and mechanism of DSF treatment on IPF, thereby realizing DSF repositioning in clinic. Together, DSF inhibited the EMT and ECM in human primary DHLF-IPF cells and TGF-β1 stimulated A549 cells via activating COX-2/PGE2/EPs axis. In vivo evidence showed that DSF significantly repressed EMT and ECM deposition via upregulated PGE2 level in BLM induced IPF mice to retard fibrosis progress, suggesting the potential anti-fibrosis effect in IPF.

## 2. Materials and methods

### 2.1. Cell culture and reagents

Human type II alveolar epithelial cells (A549) were purchased from Fenghui Biological Technology and cultured with DMEM medium contained with 10% FBS and 1% penicillin/streptomycin. DHLF-IPF cells were contributed by Professor Ren and Dr. Cao and cultured with F-12K medium contained with 10% FBS and 1% penicillin/streptomycin. Cells were cultured at 37℃ and 5% CO_2_. A549 cells retain type Ⅱ alveolar epithelial-like characteristics and can be stimulated by TGF-β1 to transform into mesenchyma, which is used for the experimental study of IPF[20].

### 2.2. Cell viability and cell death

After cultured with TGF-β1 (10 ng/mL) for 24 hours, A549 cells transformed into mesenchymal-like cell[21], then TGF-β1-induced A549 cells and DHLF-IPF cells were treated with different concentration gradient DSF for 24 hours at a density of 8×10^5^/mL. Cell Counting Kit-8 (CCK-8) and propidium iodide/crystal violet (PI/CV) were added to evaluate cell viability and cell death.

### 2.3. Wound-healing assays

TGF-β1-induced A549 cells and DHLF-IPF cells were planted at 10^5^ cells per well in a 6-well plate. Tips were used to scratch the cells in the center of well plate. Images of the scratch breadth were examined and collected using light microscopy imaging at various time points and analyzed using Image-J software.

### 2.4. Western blot

After lysed with RIPA buffer, the mixture of cells and the mice lung tissues were then centrifuged to collect the supernatant. The concentration of total protein was detected by BCA kit, and each lane of the SDS-polyacrylamide gels received equal protein. According classical western-blot, the results were analyzed by Image-J software. The antibodies were collected at below.

#### Antibodies

Anti-COL1A1 (COL-I) and anti-α-smooth muscle actin (α-SMA) were from Santa Cruz Biotechnology (#sc-293182 and #sc-53142), Santa Cruz, CA, USA. Anti-fibronectin (FN) (#610077) was from BD Biosciences, New Jersey, USA). Anti-α-smooth muscle actin (α-SMA), anti-E-cadherin (E-cad), anti-vimentin (VIM), anti-COX-2 were from Cell Signaling Technology (#19245, #14472, #5741and #12282, Danvers, MA, USA. Anti-EP1 receptor and anti-EP3 receptor were from Cayman, Michigan, USA (#101740 and #101760). Anti-GAPDH (#ab8245) was from Abcam, Cambridge, UK.

### 2.5. RNA isolation and quantitative real-time PCR

Total RNA was harvested following TRIzol Reagent manufacturer’s instructions. cDNA was obtained by reverse transcription of RNA. Fluorescent labeling was done using a SuperReal PreMix Plus (SYBR Green) and Real-time quantitative PCR was performed with the Bio-Rad CFX Maestro System. The expression of mRNA was normalized to GAPDH expression. Human and mice primer sequences were collected at below.

#### Human primer sequences

GAPDH (F 5’ TCCAAAATCAAGTGGGGC 3’, R 5’ ACTACTAGAACTCCGACA 3’),

COL1A1 (F 5’ GAGGGCCAAGACGAAGACATC 3’, R 5’ CAGATCACGTCATCGCACAAC 3’),

ACTA2(F 5’ GTGTTGCCCCTGAAGAGCAT 3’, R 5’ GCTGGGACATTGAAAGTCTCA 3 ’)

CDH1 (F 5’ CGAGAGCTACACGTTCACGG3’, R 5’ GGGTGTCGAGGGAAAAATAGG 3’)

VIM (F 5’ AGTCCACTGAGTACCGGAGAC 3’, R 5’ CATTTCACGCATCTGGCGTTC 3’)

FN1 (F 5’ CGGTGGCTGTCAGTCAAAG 3’, R 5’ AAACCTCGGCTTCCTCCATAA 3’)

PTGR1 (F 5’ AGCTTGTCGGTATCATGGTGG 3’, R 5’ AGCAAGTGTATGACCCTGGTAAT 3’)

PTGER3(F 5’ CGCCTCAACCACTCCTACAC 3’, R 5’ GACACCGATCCGCAATCCTC 3’)

PTGS2 (F 5’ CTGGCGCTCAGCCATACAG 3’, R 5’ CGCACTTATACTGGTCAAATCCC 3’)

#### Mice primer sequences

Gapdh (F 5’ CATCACTGCCACCCAGAAGACTG 3’, R 5’ ATGCCAGTGAGCTTCCCGTTCAG 3’)

Col1a1 (F 5’ GCTCCTCTTAGGGGCCACT 3’, R 5’ ATTGGGGACCCTTAGGCCAT 3’)

Acta2 (F 5’ GGCACCACTGAACCCTAAGG3’, R 5’ ACAATACCAGTTGTACGTCCAGA 3’)

Cdh1 (F 5’ TCGGAAGACTCCCGATTCAAA 3’, R 5’ CGGACGAGGAAACTGGTCTC 3)

Vim (F 5’ CCACACGCACCTACAGTCT 3’, R 5’ CCGAGGACCGGGTCACATA 3’)

Fn1 (F 5’ TCAAGTGTGATCCCCATGAAG 3’, R 5’ CAGGTCTACGGCAGTTGTCA 3’)

Ptgs2 (F 5’ TTCCAATCCATGTCAAAACCGT 3’, R 5’ AGTCCGGGTACAGTCACACTT 3’)

### 2.6. Lentiviral construction and infection in A549 cells

Three short hairpin (sh)RNA vectors targeting COX-2 (shCOX-2) and a control vector (shNC) were designed and purchased from GenePharma. Lentiviral particles were produced by transfecting HEK293T cells with lentiviral plasmids along with envelope (VSVG) and packing plasmids. For viral infection, A549 cells were plated in 6-well plates, grown to 50-70% confluence, and infected with the presence of 8 μg/Ml polybrene. Following infection for 48 hours, the cells were selected with 5.0 μg/mL puromycin. Knockdown efficiencies were confirmed via real-time PCR and western blot analysis.

### 2.7. Immunofluorescence microscopy

The cell slides were washed 3 times with PBS and then fixed with 4% paraformaldehyde for 15 min in culture plates. Cells were permeabilized with 0.5% Triton X-100 for 15 min at room temperature. Slides were dropped with 10% goat serum and blocked for 1 h at room temperature. The blocking solution was removed by absorbent paper, and diluted primary antibody was added to each slide and incubated in a wet box at 4 ℃ overnight. The primary antibody was removed by absorbent paper, followed by fluorescent secondary antibody and incubated for 1 h at 37 ℃ in a black wet box. Finally, DAPI was added and incubated in the dark for 5 min to stain nuclei. Slides were sealed with antifade solution containing anti-fluorescence quencher, and images were observed and collected under a fluorescence microscope. Pictures were analyzed with Image-J software.

### 2.8. PGE_2_ measurement

After treated with DSF, the supernatant of cells was collected after centrifuged at 4°C for 5 min at 1000 rpm/min. The remanding cells were stained with purple crystal to quantitate total protein. The PGE_2_ concentration of the supernatants and serum from mice was determined according to the manufacturer’s instructions, and the PGE_2_ concentration in supernatants normalized to the total protein.

### 2.9. BLM-induced IPF in mice

Males C57BL/6J mice weighted 20±2 g (Charles Rive) and housed at 22-24°C with a 12:12 hr light-dark cycle. Animal experiments were performed according to the Guidelines on Laboratory Animals of Nankai University and were approved by the Institute Research Ethics Committee at Nankai University (approval number: 2021-SYDWLL-000461).

The establishment and measured of IPF mice model referred to previous studies[21]. 50 mg/kg DSF was intraperitoneal injection daily for 14 days beginning 7 days after BLM administration, 0.5% CMC-Na was used as a vehicle.

### 2.10. Histology and immunohistochemistry

Before the lung tissue of mice was embedded in paraffin and sectioned, it was fixed with 4% paraformaldehyde for 2 days. Tissue paraffin sections were stained with Hematoxylin-Eosin (H&E) Staining Kit or Masson’s Trichrome Stain Kit. Tissue slices were treated with 3%-hydrogenperoxide solution to remove endogenous enzymes, infiltrated with 0.5% Triton-100 to permeabilize membrane and blocked by 10% goat serum. Slides removed the blocking solution, then added the primary antibody dropwise, and incubated overnight at 4°C. Add the secondary antibody working solution for 1 hours at room-temperature. Slides were dropped with DAB working solution and counterstained with hematoxylin. Stained tissue slices were observed under the microscope. Pictures were analyzed with Image-J software.

### 2.11. Hydroxyproline Assay

Accurately weigh the right lung and follow the instructions of the hydroxyproline test kit purchased from Nanjing Jiancheng. Results were expressed as µg of hydroxyproline/mg of lung weight.

### 2.12. Human subjects

The lung tissues and serum of IPF used in the study were provided by Professor Dunqiang Ren (Peking Union Medical College Hospital). The control lung tissues were derived from the non-tumor infiltrated area of lung cancer patients. Control serum was obtained from patients without pulmonary fibrosis. The study complied with medical ethics (Approval number: NKUIRB2021106).

### 2.13. Statistical analysis

All data were presented as the means ± SEM of at least three independent experiments (n≥3). The Student’s t test was used to compare two groups and two-way ANOVA was used for multiple group comparisons. Statistical significance was considered at *P* < 0.05. The graphical representation and statistical analysis were performed using GraphPad Prism (Version 8.3.0).

## 3. Results

### 3.1 DSF inhibited viability and migration of DHLF-IPF and TGF-β1-induced A549 cells

Cell culture models and human lung primary cells are beneficial for exploring the mechanism of EMT, lung fibrosis and the associated treatment strategies. TGF-β1 is a prototype mediator for fibroblast differentiation into myofibroblasts, induction of alveolar epithelial cells transformation into mesenchymal cells, as well as the phenotypic mediator for extracellular matrix [22]. Therefore, we stimulated alveolar epithelium A549 cells were stimulated with TGF-β1 (10ng/ml) for 24 h to establish an EMT model *in vitro*.

Next, to determine the cultured cell treatment dose of DSF, cell death and cell viability assay were performed to assess the induction of cell viability in cultures following treatment with DSF at the indicated dose. After treating with DSF for 24 h in TGF-β1-induced A549 cells and DHLF-IPF cells, cell viability and cell death were measured with CCK8 (Figures 1A and 1D) and PI/CV (Figures 1B and 1E), respectively. The half-maximal inhibitory concentrations (IC_50_) of DSF in DHLF-IPF cells and TGF-β1-induced A549 cells were 14.84 μM (Figure 1A) and 20.99 μM (Figure 1D) respectively.

**Figure 1.**
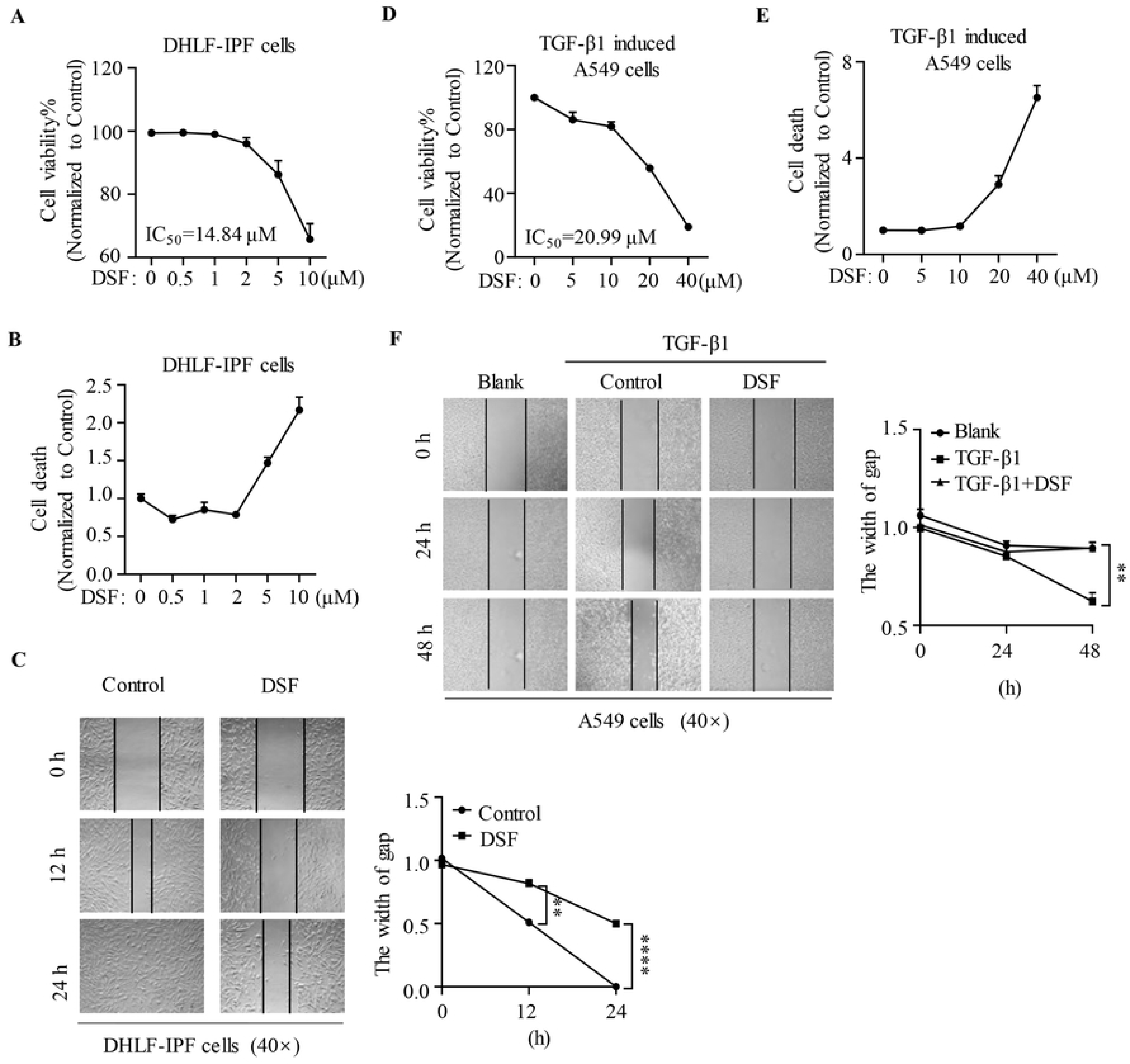
DSF inhibited viability and migration of DHLF-IPF and TGF-β1-induced A549 cells. Primary DHLF-IPF cells and TGF-β1-induced A549 cells and were exposed to indicated dose of DSF for 24 h. Cell viability **(A and D)** and cell death **(B and E)** were determined by a CCK-8 staining assay and PI exclusion assay, respectively. The half-maximal inhibitory concentration (IC_50_) was calculated by cell viability **(A and D)**. The width of the scratch was photographed and quantified at 0, 12, and 24 h post scratching of DHLF-IPF cells **(C)** or at 0, 24, and 48 h post scratching of TGF-β1-induced A549 cells **(F)** (magnification 40×) by a wound-healing assay. The width of gap was measured with Image-J software (Three independent analyses were performed) and calculated with GraphPad Prism. \**P*<0.05, \*\**P*<0.01, \*\*\*\**P*<0.0001.

Both DHLF-IPF cells and TGF-β1-induced A549 cells showed dose and time-dependent responses to DSF treatment. In a following antifibrosis study, we used 5 μM DSFfor DHLF-IPF cells and 15 μM for TGF-β1-induced A549 cells to avoid interference from cytocidal effects according to IC_50_.

TGF-β1-induced A549 cells were characterized with EMT phenotype and generated a migratory phenotype. Thereby, we evaluated the effect of DSF on cell migration via an *in vitro* wound healing assay, and results revealed that cell migration rates were significantly reduced in both primary DHLF-IPF cells (Figures 1C) and TGF-β1-induced A549 cells (Figures 1F). Together, these results suggested that DSF inhibited cell viability in a dose-dependent manner, accompanied with cell migration impeded during EMT progress.

### 3.2 DSF reversed EMT and ECM in DHLF-IPF and TGF-β1-induced A549 cells

Since the unexpected wound-healing capacities seen in the context of DSF, the effect of regulatory effects of DSF on EMT and ECM-related biomarkers in DHLF-IPF cells and TGF-β1-induced A549 cells were further investigated.

DSF (5 μM) depressed the mRNA expression of mesenchymal markers *CDH2*, *VIM* and *ACTA2,* as well as extracellular matrix *COL1A1* (Figure 2A), as well as the protein levels of VIM, α-SMA and FN in DHLF-IPF cells (Figures 2B **and Supplementary Figure 1A**). Accordingly, DSF (15 μM) were added to TGF-β1-induced A549 cells in the presence of TGF-β1 for 24 h, the mRNA level of epithelial marker *CDH1* was increased, *CDH2* and *COL1A1* were reduced significantly compared with TGF-β1 group (Figure 2C). Similarly, the protein expression of α-SMA, VIM and FN were depressed whereas the epithelial marker E-cad was not significantly increased (Figures 2D **Supplementary Figure 1B**). What’s more, western blot analysis was supported by immunofluorescence results showing a significant decrease in cellular VIM expression occurred after 24 h treatment of DSF in TGF-β1-induced A549 cells (Figure 2E). The reverse change in mesenchymal proteins and epithelial marker, as well as the reduced ECM deposition suggested that the process of EMT was disrupted by DSF.

**Figure 2.**
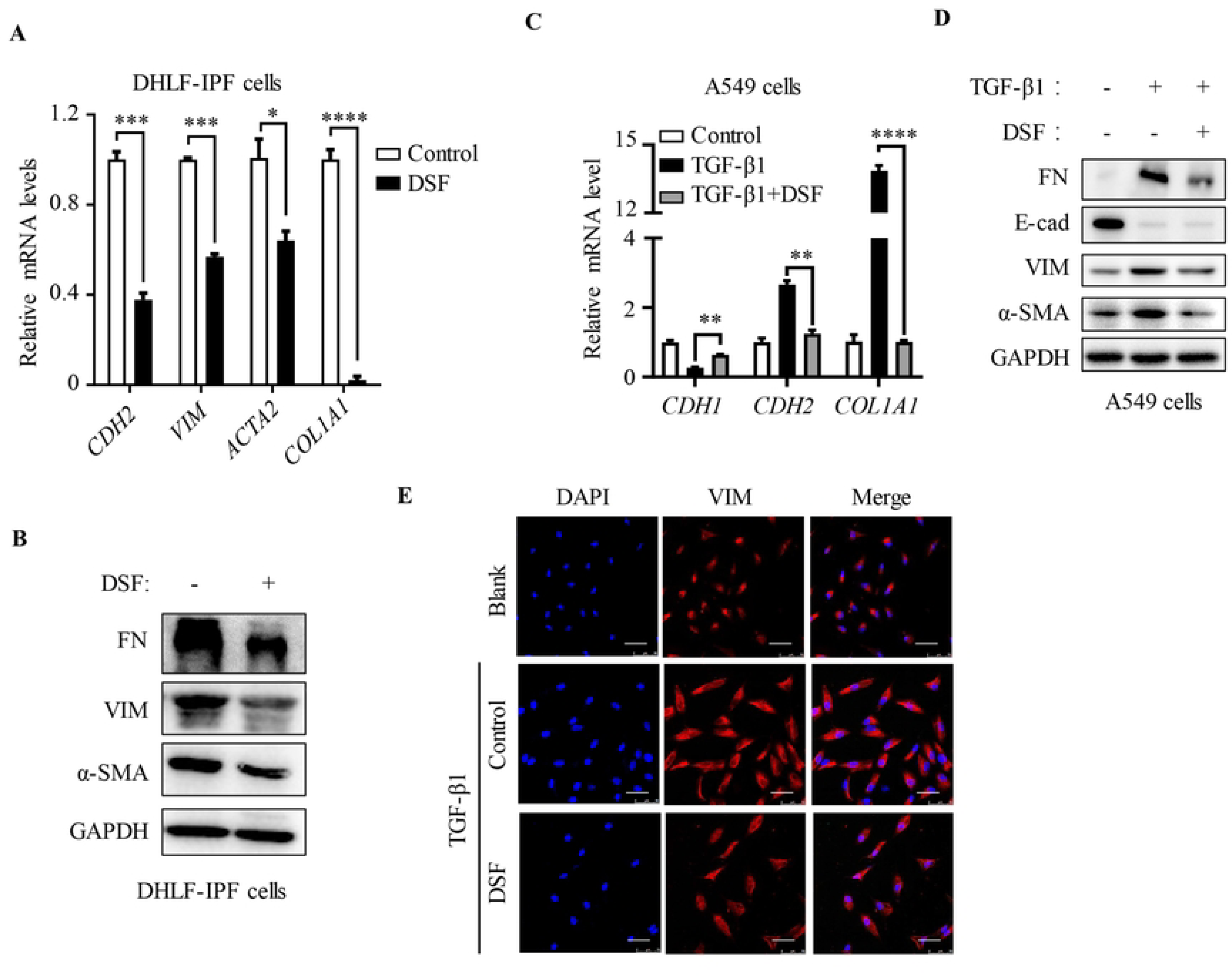
DSF reversed EMT and ECM in DHLF-IPF and TGF-β1-induced A549 cells. **(A)** The mRNA levels in DHLF-IPF cells including *CDH2*, *VIM*, *ACTA2* and *COL1A1* were detected by qPCR. **(B)** The protein expression of VIM, α-SMA and FN were measured with western blot in DHLF-IPF cells. After induced with or without 10 ng/ml TGF-β1 for 24 h, TGF-β1-induced A549 cells were treated with DSF (15 μM) for another 24 h. **(C)** mRNA levels of *CDH1* and *CDH2* and *COL1A1* were detected by qPCR. **(D)** The protein expression of E-cad, VIM, α-SMA and FN were measured with western blot. **(E)** Immunofluorescence staining of VIM were performed and nuclear staining with DAPI in TGF-β1-induced A549 cells (magnification 400×, bar=50 μm). \**P* <0.05, \*\**P*<0.01, \*\*\*\**P*<0.0001.

### 3.3 DSF inhibited TGF-β1-induced EMT through restoring COX-2 regulated PGE_2_ biosynthesis

During IPF development and progress, α-SMA is considered as a gold standard and regarded as a marker of active fibrogenesis [23]. Firstly, the disordered structure (H&E), significant fibrosis (Masson’s staining), α-SMA and FN positive expression in the lung tissues was observed compared with those in non-IPF through the histological alterations in human lung tissues with IPF (**Supplementary Figure 2A**), suggesting the EMT progress and ECM deposition. We then reanalyzed a public dataset (GEO accession #: GSE10667), and found that *PTGS2* mRNA was significantly reduced in IPF lung tissues (Figure 3A), which made us curious about the relationship between the COX-2 and EMT, thus we measured the differences of COX-2 and α-SMA in comparable regions of lung tissue from IPF patients using immunohistochemistry (Figure 3B) and confocal microscopy (Figure 3C). Immunohistochemistry was used to examine the spatial location of COX-2 and α-SMA in lung tissues from IPF patients (Figure 3B). In case #1 IPF lung tissue (left), it showed that low COX-2 expression located in a α-SMA-positive tissue area. On the contrary, case #2 showed high positive of COX-2 and lack of α-SMA expression (Figure 3B), demonstrating the potential negative relationship between the expression of COX-2 and α-SMA. Furthermore, limited co-localization between COX-2 and α-SMA was present in IPF patients via immunofluorescence microscopy (Figure 3C). In addition, COX-2 metabolite PGE_2_ production in serum was detected via Elisa assay, and result revealed that it was decreased isolated in serum from IPF patients compared with healthy donors (Figure 3D), suggesting COX-2/PGE_2_ axis may play an essential role in IPF development and progression.

**Figure 3.**
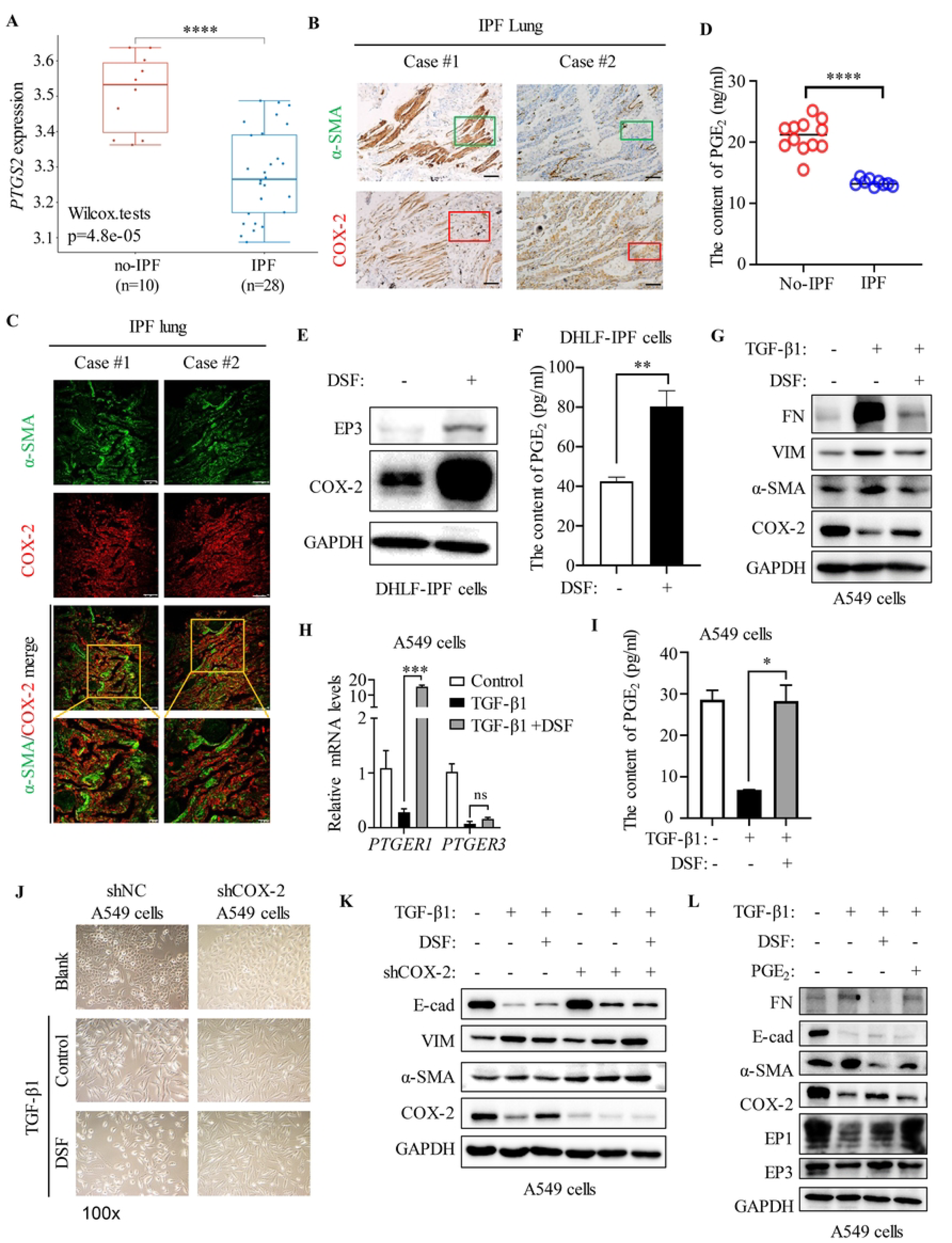
DSF inhibited TGF-β1-induced EMT through restoring COX-2 regulated PGE2 biosynthesis. **(A)** Reanalyzed of *PTGS2* mRNA from a public dataset (GEO accession #: GSE10667) with Wilcox. tests. **(B)** The images showed the expression of COX-2 and α-SMA staining in IPF patient tissues. α-SMA staining (green frames) and COX-2 expression (red frames) were observed in fibroblastic foci (magnification 200×). **(C)** Immunofluorescence analysis showed the expression of α-SMA (green) and COX-2 (red) staining in IPF patient lung tissues. IPF lung tissues area positived for COX-2 presented α-SMA negativity (scale bar=75 μm or 25 μm). **(D)** The prostaglandin E2 (PGE_2_) content from no-IPF (n=12) and IPF (n=9) patient serum was assayed by ELISA kit. **(E)** DHLF-IPF cells were treated with DSF (5 μM) for 24 h, then the protein expression of COX-2 and EP3 were measured with western blot. **(F)** The PGE_2_ content in the supernatant was measured by ELISA kit after DHLF-IPF cells were treated with DSF (5 μM) for 24 h. After induced with or without 10 ng/mL TGF-β1 for 24 h, TGF-β1-induced A549 cells were treated with DSF (15 μM) for another 24 h. **(G)** The protein expression of COX-2, α-SMA, VIM and FN and were measured with western blot in A549 cells. **(H)** The mRNA levels of PGE_2_ receptors (*PTGER1* and *PTGER3*) were detected by qPCR and normalized with *GAPDH* in A549 cells. **(I)** TGF-β1-induced A549 cells were treated with DSF (15 μM) for 24 h, then the prostaglandin E2 (PGE_2_) content in the supernatant was assayed by ELISA kit. **(J)** Cell morphology changes were observed and photographed with a light microscopy (magnification 100×). **(K)** Both shCOX-2 and shNC A549 cells were treated with DSF (15 μM) for 24h, then the protein levels of E-cad, VIM, α-SMA and COX-2 were assessed with western blot. **(L)** TGF-β1-induced A549 cells were treated with DSF (15 μM) or PGE_2_ (5 μM) for 24 h. The protein expression was measured with western blot. \**P* < 0.05, \*\**P* < 0.01,\*\*\**P* <0.001, \*\*\*\**P*<0.0001.

COX-2 is the rate-limiting enzyme in the metabolic conversion of arachidonic acid (AA) into various prostaglandins (PGs) including prostaglandin E2 (PGE_2_) [24]. Although some studies showed that PGE_2_ had pro-inflammatory actions, accumulating data suggested that the COX-2/PGE_2_ plays a vital role in ameliorating fibrosis and avoiding respiratory damage in IPF [25].

To further confirm our conjecture, COX-2 inhibitors Rofecoxib was performed in our following experiments. The administration of COX-2 inhibitors Rofecoxib did promote EMT through re-expression of VIM, α-SMA and FN in IPF cells (**Supplementary Figure 3B and 3C**), as well as the depressing PGE_2_ production in A549 cells (**Supplementary Figure 3D**), suggesting the loss of COX-2 promoted EMT. In view of diminished COX-2 expression in fibroblasts with a resultant defect in the antifibrotic mediator PGE_2_ production in IPF, we tested whether DSF treated IPF through activating COX-2 to induce PGE_2_ production. We treated DHLF-IPF cells with DSF (5 μM), and detected relevant indicators through western blot and Elisa. Results exactly suggested that the level of COX-2, PGE_2_ receptor-3 (EP3) (Figure 3E **and Supplementary Figure 3A**) and PGE_2_ content (Figure 3F) in supernatant was increased in primary DHLF-IPF cells with DSF treatment. Likewise, DSF induced COX-2 (Figure 3G **and Supplementary Figure 3E**) expression and PGE_2_ receptors (PTGER1 and PTGER3) (Figure 3H), which increased prostaglandin E2 (PGE_2_) level (Figure 3I), and subsequently improved EMT through the downregulation of α-SMA, VIM and FN (Figure 3G **and Supplementary Figure 3E**) in A549 cells.

Given the significant COX-2 expression difference and relevance of IPF, we further evaluated the direct roles of COX-2 in IPF. Then, COX-2-targeting shRNA (shCOX-2) or corresponding controls (shNC) were used to establish a stable COX-2-knockdown cell line in A549 cells (**Supplementary Figure 3F**). Cell morphology and protein results revealed that DSF had limited interference on TGF-β1 induced shCOX-2 A549 cells, which showed no significant changes in cell migration morphology (Figure 3J) and the expression of EMT and ECM markers (Figure 3K **and Supplementary Figure 3G**) compared with corresponding shNC cells. To sum up, these conclusions suggested that DSF may mediate COX-2 expression to play its role in the treatment of IPF.

After determining the role of COX-2, we continue to explore the function of its downstream product PGE_2_ in IPF. Unsurprisingly, TGF-β1 inhibited EMT and COX-2/PGE2 signaling pathway, while DSF treatment reversed this phenomenon. Likewise, exogenous PGE_2_ (5 μM) treatment for 24 h in the presence of TGF-β1 activated EP1 and reversed the expression of VIM, α-SMA and FN, though did not increased COX-2 expression in TGF-β1 induced A549 cells compared with DSF treatment (Figure 3L **and Supplementary Figure 3H**). Together these data strongly suggested that the expression of COX-2 made important contribution to the pathogenesis of pulmonary fibrosis. DSF is associated with upregulation of COX-2, which in turn promotes PGE_2_ synthesis and secretion to improve EMT and ECM.

### 3.4 Anti-fibrotic effect of DSF by boosting PGE2 biosynthesis in BLM-induced IPF model

Finally, experimental models of fibrosis *in vivo* are available for defining the complexity of matrix metabolism in the intact tissue and validating the findings from cell culture and *in vitro* systems. IPF mice model was established using 2mg/ml BLM by atomized drug delivery device for 7days. Mice were sacrificed at the endpoint after DSF treatment for 14 days. BLM mice treated with or without DSF showed limited differences, but their body weights were lower than those of blank mice (Figure 4A). For pulmonary function, DSF (50 mg/kg) treatment attractively relieved respiratory system dysfunction in the preclinical model via enhancing FVC, Cdyn and depressing Re, Ri compared with BLM treated group (Figure 4B).

**Figure 4.**
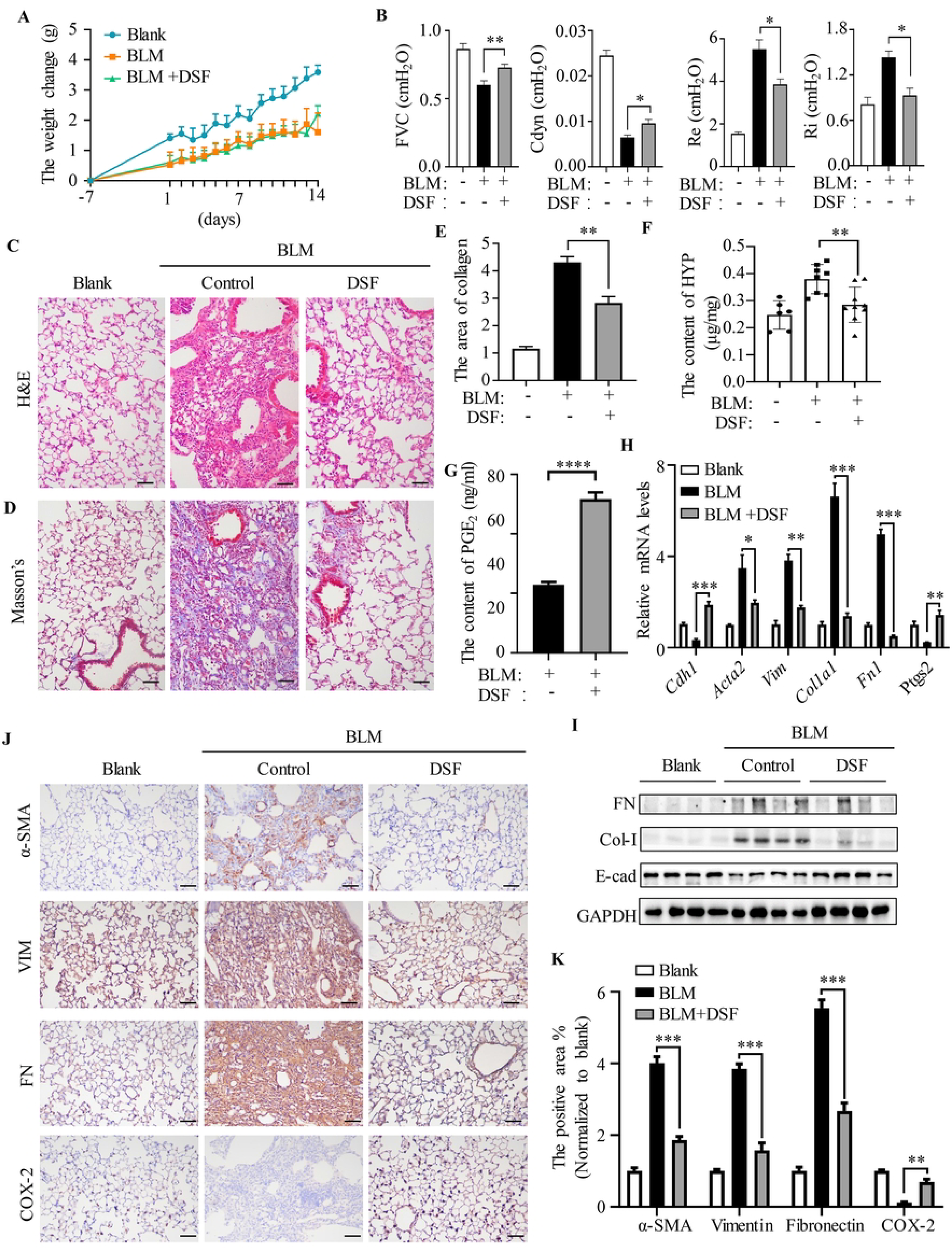
Anti-fibrotic effect of DSF by boosting PGE2 biosynthesis in BLM-induced IPF model. **(A)** The body weight of each mouse was monitored and recorded daily. **(B)** Pulmonary function paraments including forced vital capacity (FVC), dynamic compliance (Cdyn), expiratory resistance (Re) and inspiratory resistance (Ri) among different treatments were measured after treated with DSF for 14 days. Lung sections were stained with H&E **(C)** or Masson’s trichrome **(D)** for collagen accumulation (representative image, magnification 200×, bar=100 μm), and Masson’s trichrome staining was quantified **(E)** by Image-J software compared to blank group. **(F)** The hydroxyproline content in lung tissues among different groups were analyzed and quantified. **(G)** The prostaglandin E2 (PGE_2_) content in the mice serum were detected by ELISA kit. **(H)** The mRNA levels of *Cdh1, Acta2, Vim, Col1a1, Fn1* and *Ptgs2* in lung tissues were performed by qPCR and normalized with *Gapdh*. **(I)** Western blot was used to analyze the expression of E-cad, FN and Col-Ⅰ in lung tissues. **(J)** Immunohistochemistry staining of FN, VIM and α-SMA and COX-2 in the lung tissues (magnification 200 ×, scale bar = 100 μm). **(K)** The positive area on lung sections was quantified by Image-J software, normalized to blank control. \**P*<0.05, \*\**P*<0.01,\*\*\**P*<0.001, \*\*\*\**P*<0.0001.

H&E staining was used to observe changes in mice lung tissue pathological structure. The alveolar structure in the BLM group was blurred or even disappeared, and the alveolar shape was incomplete combined with obvious fibrotic foci. Meanwhile, the cell nucleus was deeply stained and the cell proliferation increased wildly. On the contrary, the most of the intact alveolar structure of BLM-mice treated with DSF (50 mg/kg) was retained, and the fibrotic foci were reduced significantly with no obvious cell proliferation (Figure 4C). Masson’s trichrome staining further suggested the collagen deposition in lung tissues, which showed a significant increase of collagen around the fibrotic foci in BLM-mice lung tissue compared with the control mice (Figure 5D and 5E). In addition, collagen deposition in lung sections also quantified from the hydroxyproline content (Figure 5F), both were strikingly decreased with DSF (50 mg/kg) treatment in fibrotic foci of BLM mice compared with those in the control group. At the end point of the experiment, mice lung tissues were removed for further tissue proteins, mRNA and IHC staining analysis to evaluate the effect of DSF on BLM-induced IPF. DSF (50 mg/kg) reduced the mRNA levels of mesenchymal markers (*Vim*, *Acta2*), ECM markers (*Fn1*, *Col1a1*) and increased epithelial marker (*Cdh1*) compared to the BLM group (Figure 4H). Simultaneously, the protein levels in lysates of whole lung tissues were analyzed, and DSF (50 mg/kg) treatment effectively suppressed Col-Ⅰ and FN expression and increased E-cad expression (Figure 4I **and Supplementary Figure 4A**). Similarly, immunohistochemistry results further showed that BLM induced the level of FN, VIM and α-SMA in mice as compared with control mice, whereas DSF (50 mg/kg) treatment significantly reduced BLM-induced the overexpression of FN, VIM and α-SMA expression (Figure 5J and 5K).

**Figure 5.**
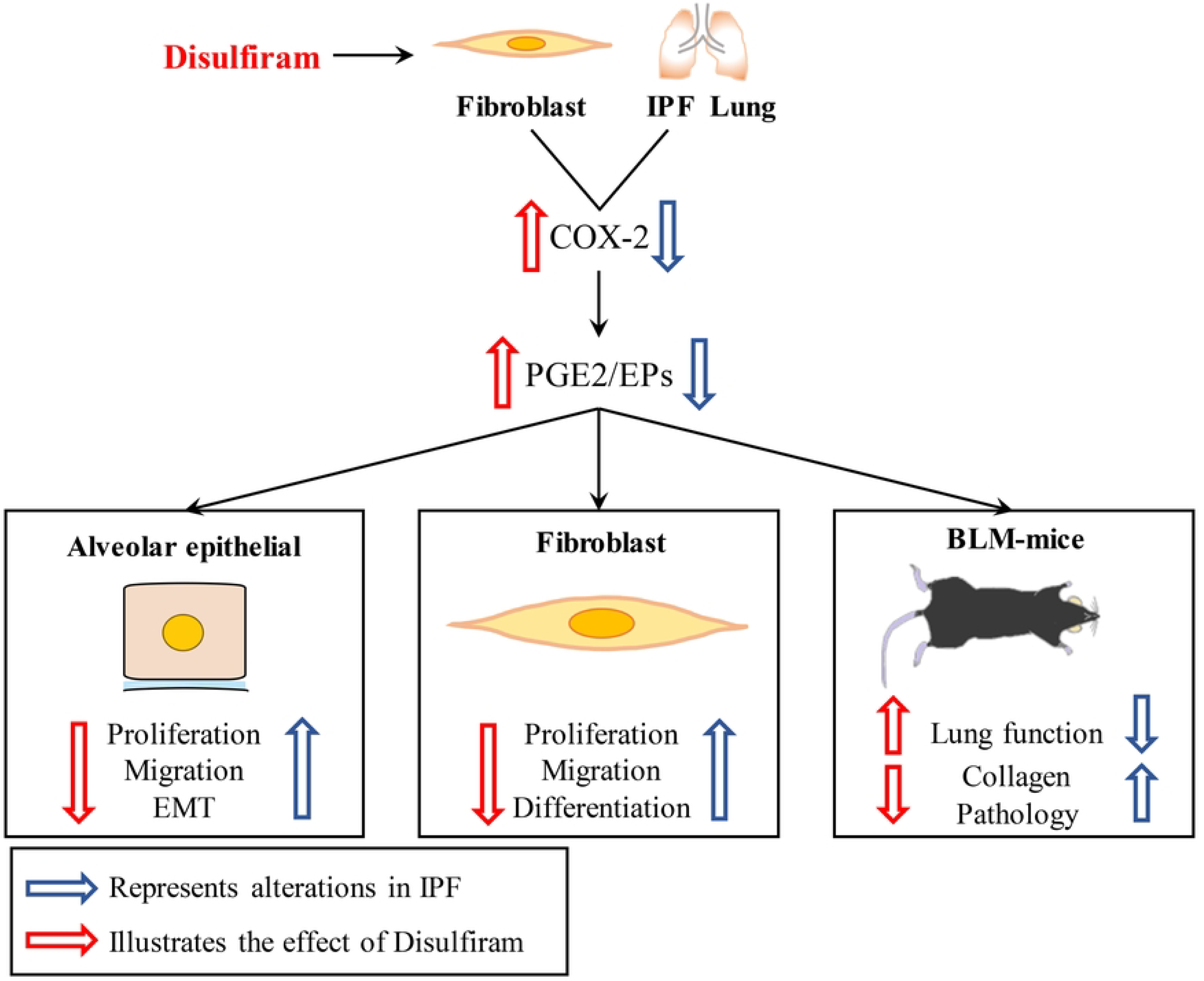
Scheme. We find that COX-2/PGE_2_ is negatively expressed in IPF patients. The decrease of COX-2 promotes abnormal cell proliferation, induces the epithelial-mesenchymal transition (EMT) of alveolar epithelial cells, activates fibroblast differentiation, and reduces the production of collagen. Whereas, disulfiram exerts the effect of inhibiting cell proliferation and migration, decreasing EMT of alveolar epithelial cells, as well as preventing fibroblast activation. On the other hand, DSF also ameliorates lung function, collagen deposition and pathology injure in BLM induced IPF mice.

Further we explored the mechanism of DSF *in vivo*, we confirmed the mRNA level of *Ptgs2* (Figure 5H) and the protein of COX-2 positive expression (Figure 5J and 5K) in whole lung tissue and lung section respectively *in vivo* experimental IPF mice. In addition, the content of PGE_2_ in mice serum was increased in DSF group compared with BLM-only group (Figure 5G). These data indicated that DSF (50 mg/kg) treatment significantly reduced BLM-induced EMT progression and ECM deposition *in vivo*, accompanied with pulmonary function reparation and COX-2 reactivation to mediate PGE_2_ biosynthesis, thus ameliorating IFP progression.

## 4. Discussion

This study revealed the anti-IPF pharmacological activity of the anti-alcohol abuse drug disulfiram (DSF) [26] that rarely explored. In our research, we utilized human primary DHLF-IPF cells and TGF-β1 induced EMT cells as the *in vitro* model. Besides, intratracheal injection of BLM into mice induced IPF mice model to estimate the anti-fibrotic effect of DSF *in vivo*. Our results proved that DSF inhibited the proliferation and migration in IPF cell model, improved IPF mice respiratory function and prevented lung fibrosis. Meanwhile, DSF increased epithelial proteins, reduced mesenchymal proteins and excessively deposited extracellular matrix proteins *in vitro* and *in vivo*. Notably, DSF regulated EMT by activating PGE_2_ biosynthesis, and the anti-IPF pharmacological activity of DSF have not reported so far.

The formation mechanisms of IPF mainly include the transformation of alveolar epithelial cells to mesenchyme [27], activation of myofibroblasts [28], deposition of extracellular fibrous protein [29], secretion of cytokines [30] and so on. TGF-β1 is a recognized pathogenic factor for pulmonary fibrosis [31]. We have demonstrated for the first time that DSF exhibited admirable effect on improving EMT and degrading extracellular matrix protein on TGF-β1 induced pulmonary fibrosis cell models and DHLF-IPF cells.

Previous studies confirmed that the expression of COX-2 and PGE_2_ was down-regulated in myofibroblasts and IPF patients [8], while α-SMA is highly expressed in lung fibrous foci [8]. Likewise, our analysis of lung pathology in IPF patients also found that α-SMA and COX-2 were not co-localized and the expression of PGE_2_ was decreased in the serum. These results proved that COX-2/PGE_2_ was a possibility target for IPF.

Numerous studies showed that TGF-β1 induced COX-2 and PGE_2_ expression [32]. TGF-β1 induced the expression of COX-2 and increased the synthesis of PGE_2_ in prostate cancer cells [33]. TGF-β1 induced COX-2 expression to train EMT in human bronchial epithelial cells [34]. TGF-β1 increased COX-2 and PGE_2_ receptor EP2 expression in breast cancer cells [35], and supported that PGE_2_ was a mediator to incite angiogenesis and cell migration, and selective EP2 inhibitors reduced the expression of PGE_2_ [35]. Conversely, Peedikayil E Thomas et.al explained that PGE_2_ showed significant effect on inhibiting TGF-β1 induced myofibroblast differentiation, including modulating cell morphology, cytoskeleton, and cell adhesion-dependent signals [36]. In addition, transcriptome analysis of TGF-β1 induced myofibroblasts differentiation process found that PGE_2_ reversed the expression of 363 (62%) TGF-β1 up-regulated genes and 345 (50%) TGF-β1 down-regulated genes [37]. Our results revealed that TGF-β1 reduced COX-2 and PGE_2_ expression, and COX-2 silence A549 cells are more susceptible to TGF-β1, thus aggravating EMT development. We observed that exogenous addition of PGE_2_ improved EMT and ECM induced by TGF-β1. These results indicated the important role of COX-2/PGE_2_ in IPF.

In recent years, the application research of DSF has been ever more extensive [38]. DSF alone or chelated with divalent metal ions exerted anti-cancer activity [39]. In addition, DSF was realized as a narrow-spectrum antibacterial agent [40, 41]. DSF dose-dependently inhibited the level of PGE_2_ and COX-2 protein expression in the aqueous humor of uveitis rats whatever oral [42] or topical eye medication [43]. DSF eye drops administration inhibited the deposition of fibrotic protein in ocular scar formation in mice. Mechanically, DSF mainly suppressed inflammation factors to improve fibrous lesions [15]. Studies have shown that DSF inhibits the secretion of inflammatory factors and type Ⅰ collagen in rat unilateral urethral obstruction model [44]. What’s more, the main metabolite of DSF, diethyldithiocarbamate (DDC), suppressed the inflammation and fibrosis-related parameters in non-alcoholic fatty liver by regulating lipid metabolism and oxidative stress in rodents, including the inhibition of collagen deposition and expression of α-SMA protein in liver [45]. PGE_2_ often served as an effective pro-inflammatory mediator and participated in the inflammatory diseases [46]. The above studies proved that DSF inhibited the inflammatory factors PGE_2_ and COX2 protein. On the contrary, we verified DSF increased COX-2 and PGE_2_ in EMT cells induced by TGF-β1, human primary DHLF-IPF cells, and IPF mice. To determine the role of COX-2 in the treatment of IPF with DSF, shCOX-2-A549 cells were induced EMT with TGF-β1 and processed by DSF. We found that DSF failed in improving the EMT and ECM parameters in the shCOX-2 EMT cell model. Instead, it played an anti-fibrotic effect by inducing the expression of COX-2. PGE_2_ is the main production mediated and catalyzed via COX-2 [47]. We concluded that exogenous addition of PGE_2_ significantly improved EMT model of TGF-β1 induced IPF. Therefore, we believed that DSF prevented EMT and treated IPF by inducing COX-2/PGE_2_ axis expression.

The actual strategy to increase PGE_2_ in lung tissue during IPF was limited. The inhibitor of 15-prostaprostaglandin dehydrogenase (15-PGDH), the PGE_2_ degrading enzyme, indirectly increased PGE_2_ content, thereby destroying TGF-β signaling and inhibiting myofibroblasts growth and differentiation [48]. In order to reduce the adverse side effects of elevated PGE_2_ on other organs, I Ivanova V et al. employed liposomes to deliver PGE_2_ into the lungs by inhalation to treat pulmonary fibrosis [49]. Nonetheless, this study [50] emphasized that IPF was an interspecific lung disease (ILD), and PGE_2_ was significantly elevated in ILD patients. It is pointed out that the COX-2/PGE_2_ axis has dual functions. On the one hand, activation of COX-2/PGE_2_ axis aggravated IPF induced by streptococcus pneumonia, but on the other hand, it also exists therapeutic effect on non-malignant IPF [50]. Our research based on experimental IPF induced by TGF-β1 and BLM. DSF mobilized COX-2/PGE_2_ axis and exhibited excellent anti-IPF effect. Furtherly, it is necessary to explore the anti-fibrosis effect of DSF in different IPF classification, and the role of COX-2/PGE_2_ induced by DSF in systemic organs.

From a broader perspective, our research illustrated the potential of drug repositioning, provided new mechanism insights, and determined new IPF treatment target and clinical trial inspiration. DSF, an old, safe and public domain drug may help save IPF patients worldwide.

## Author contributions

Xiaolin Pei, Fangxu Zheng, Yin Li, Zhoujun Lin, designed and performed *in vitro* studies, Xiaolin Pei, Fangxu Zheng, Yin Li, Tianjiao Li designed, performed and analyzed *in vivo* experiments. Xiaolin Pei, Xiao Han and Ya Feng performed and analyzed in flow cytometry experiments. Xiao Han, Zhoujun Lin, Fei Li and Juan Yang provided guidance on data processing and writing. Zhenhuan Tian contributed to the clinical samples collection. Dunqiang Ren contributed to the primary DHLF-IPF cells isolation & culture, and histopathological analysis. Xiaolin Pei, Ke Cao and Chenggang Li contributed to the study design, supervision of the study, draft and review the manuscript. All authors had full access to the data, and approved the final version of the manuscript.

## Ethics approval and consent to participate

Animal experiments were performed according to the Guidelines on Laboratory Animals of Nankai University and were approved by the Institute Research Ethics Committee at Nankai University.

## Consent for publication

Not applicable.

## Availability of data and materials

All data generated or analyzed during this study are included in this published article and are available from the corresponding author on reasonable request.

